# Characterisation of the phase-variable autotransporter Lav reveals a role in host cell adherence and biofilm formation in Non-Typeable *Haemophilus influenzae*

**DOI:** 10.1101/2021.10.15.464623

**Authors:** Zachary N. Phillips, Preeti Garai, Greg Tram, Asma-Ul Husna, Megan Staples, Keith Grimwood, Amy V. Jennison, Michael P. Jennings, Kenneth L. Brockman, John M. Atack

## Abstract

Lav is an autotransporter protein found in pathogenic *Haemophilus* and *Neisseria* species. Lav in non-typeable *Haemophilus influenzae* (NTHi) is phase-variable: the gene reversibly switches ON-OFF via changes in length of a locus-located GCAA_(n)_ simple DNA sequence repeat tract. The expression status of *lav* was examined in carriage and invasive collections of NTHi, where it was predominantly not expressed (OFF). Phenotypic study showed *lav* expression (ON) results in increased adherence to host cells, and denser biofilm formation. A survey of *Haemophilus* spp. genome sequences showed *lav* is present in ∼60% of NTHi strains, but *lav* is not present in most typeable *H. influenzae*. Sequence analysis revealed a total of five distinct variants of the Lav passenger domain present in *Haemophilus* spp., with these five variants showing a distinct lineage distribution. Determining the role of Lav in NTHi will help understand the role of this protein during distinct pathologies.

## Introduction

Non-typeable *Haemophilus influenzae* (NTHi) is a bacterial pathogen of global importance. NTHi colonizes the human nasopharynx, but is an important pathogen in middle ear infection (otitis media) in children (1), exacerbations in bacterial bronchitis, chronic obstructive pulmonary disease and bronchiectasis (2, 3), and community-acquired pneumonia, in adults (4). NTHi also causes invasive infections, and these are fatal in ∼10% of children <1 year, and in ∼25% of adults aged ≥80-years (5-7). Frequency of disease caused by NTHi is increasing annually, exacerbated by both the absence of an NTHi vaccine, and by emerging antibiotic-resistance (8). Understanding pathobiology, and identifying the stably expressed antigenic repertoire, of NTHi is crucial for the rational design of a protein subunit vaccine, but is complicated by factors like variable gene expression and low sequence conservation.

Several host-adapted bacterial pathogens are able to randomly and reversibly switch gene expression, a process known as phase-variation (9-11). Many bacterial genes phase-vary by changes in length of locus-located simple DNA sequence repeat (SSR) tracts. When SSR tracts are located in the open reading frame of a gene, this variation in length results in ON-OFF switching of expression. Phase-variable genes typically encode surface proteins such as iron acquisition factors (11), lipooligosaccharide biosynthetic enzymes (12), and adhesins (13). Phase-variation of bacterial surface features generates sub-populations of phenotypic variants, some of which may be better adapted to a particular niche, or equipped to avoid an immune response. Many bacterial surface proteins are classified as autotransporters, and these contain a C-terminal β-barrel translocator domain in the outer membrane, and an extracellular passenger domain (14). Many virulence-associated autotransporters are phase-variably expressed, including UpaE in uropathogenic *Escherichia coli* (15), Hap (16) and Hia (17) in *Haemophilus influenzae*, and NalP (18, 19), AutA (20) and AutB (21) in *Neisseria* spp. A homologue of AutB, named Lav, has been described in multiple *Haemophilus* spp. (21, 22). The *lav* gene has also been reported to be phase-variable, as a GCAA_(n)_ SSR tract is present in the *lav* open reading frame (22). Investigation into AutB in *N. meningitidis* found the protein played a role in biofilm formation, and was phase-varied OFF in available genomes (21). Study of another Lav homologue, Las, in *H. influenzae* biogroup *aegyptius*, has suggested a role in inflammatory cytokine production (23), and increased expression associated with disease progression (24). The function of Lav in NTHi has not been studied in detail, although over multiple rounds of infection the *lav* gene was shown to phase-vary OFF (25), implying selection against Lav during chronic/recurrent infections. Therefore, we sought to undertake a phenotypic characterisation of the role of Lav in NTHi, and to determine the prevalence and diversity of this protein in *Haemophilus* spp.

## Materials & Methods

### Bacterial Isolate Collections

Nasal (carriage) control samples were taken from the ORChID collection, a prospective birth cohort study of infants in South East Queensland. As part of this collection, respiratory disease symptoms were recorded daily, and weekly nasal swabs were collected, from 158 infants during their first two years of life (2010-2012) (26). All samples used as carriage controls were randomly selected from infants demonstrating no overt symptoms of respiratory illnesses either 2 weeks before or after sampling (27). Invasive NTHi isolates used for this study were isolated from patients suffering from *H. influenzae* infections in SE Queensland over a 15-year period (2001-2015) (28). Information on age, sample site, and geographical location were collected, but not on comorbidities (28).

### Bacterial growth and media

NTHi isolates were grown in brain heart infusion (BHI; Oxoid) supplemented (sBHI) with hemin (1%) and β-NAD (2 μg/ml) at 37°C in an atmosphere containing 5% (v/v) CO_2_. *Escherichia coli* strains were grown in Luria-Bertani (LB) broth, or on LB agar (LB broth +1.5% [w/v] bacteriological agar). LB was supplemented with ampicillin (100µg/mL) as required.

### SSR tract PCR and fragment analysis

Bacterial genomic DNA from invasive isolates was prepared as described previously (29). Standard methods were used throughout for PCR using GoTaq Flexi DNA polymerase according to manufacturer’s instructions (Promega), and fragment analysis was carried out as previously described (30). *lav* ON/OFF status was determined from the number of GCAA repeats in the SSR tract present in the gene (based on amplicon peak size), using Lav_F (FAM-GCCCCATTTATTTTTACTTGACAAAGG) and Lav_R (GCTCATTTGTTAATTTAGAATTGTCATAAG) primers by sizing and quantifying using the GeneScan system (Applied Biosystems International) at the Australian Genome Research Facility (AGRF; Brisbane, Australia), and traces analysed using PeakScanner software 2.0 (Applied Biosystems International). Enriched ON and OFF variants in strain 86-028NP were generated by colony screening and enrichment for GCAA tract lengths in the *lav* SSR tract.

### Cloning Lav protein fragment for generation of antisera

The Lav passenger domain and flanking region, comprising residues 250-540 of the full protein, was expressed by cloning the encoding DNA into the pET15b vector, in-frame with the N-terminal His-tag, The coding region was amplified from strain 86-028NP using primers Lav_bind-F (AGTCAGCATATGCAAGATAACTCACACGTTATCG) and Lav_bind-R (CTGACTGGATCCTTAGTGGCGGAAGCGTTGATATTG) with KOD HotStart proofreading DNA polymerase (Novagen) and cloned into the NdeI and BamHI sites of pET15b, to generate vector pET15b::Lav-bind. Expression was carried out using *E. coli* BL21 (DE3) containing the pET15b::Lav-bind vector in LB broth induced with 1mM Isopropyl β-d-1-thiogalactopyranoside (IPTG) at 37°C with shaking for 16 hours. Purification with TALON Metal Affinity Resin (Takara) was carried out from the insoluble fraction by using multiple rounds of sonication and washes in PBS containing 0.1% Tween (v/v). Following purification, pure Lav-bind was dialysed at 4°C for 12 hours in PBS, twice.

### Western Blotting

Protein lysates of whole NTHi were prepared by heating whole cell suspensions at 99°C for 40 mins. These were electrophoresed on 4-12% Bis-Tris polyacrylamide gels (Invitrogen) at 150V for 45mins in Bolt MOPS Running Buffer (Invitrogen). Samples were transferred to nitrocellulose membrane at 15V for 1h. Membranes were blocked with 5% (w/v) skim milk in Tris-buffered saline with 0.1% Tween-20 (TBS-T) by shaking overnight at 4°C. Primary mouse antibodies against the Lav-bind protein (anti-Lav antisera) were raised in BALB/c mice at the Institute for Glycomics Animal Facility. 50µg of purified Lav-bind protein in alum was used per mouse. Primary antibody was used at 1:1000 dilution in 5% (w/v) skim milk in TBS-T for 1h with shaking at room temperature. Membranes were washed multiple times in TBS-T for 1h before adding secondary antibody (goat anti-mouse alkaline phosphatase conjugate; Sigma) as above at 1:2500 dilution. Membranes were washed for 1h in TBS-T, before developing at room temperature with SigmaFAST BCIP/NBT prepared according to manufacturer’s instructions (Sigma).

### Animal Ethics

Animal work was approved by Griffith University Animal Ethics Committee Protocol Number GLY/16/19/AEC. Animals were cared for and handled in accordance with the guidelines of the Australian National Health and Medical Research Council (NHMRC).

### Adherence and invasion assays with host cells

NTHi adherence and invasion was assessed as previously (31, 32). Approximately 2.5 × 10^5^ A549 cells were seeded into each well of a flat-bottomed 24 well plate (Greiner, Germany) and allowed to settle overnight (37°C) before inoculating with NTHi at an MOI of 30:1, or 8×10^6^ CFU in 250µL of RPMI media (Dubco) containing 10% (v/v) foetal calf serum (FCS). Plates were incubated for 4h at 37°C with 5% (v/v) CO_2_. Wells were washed of non-adherent NTHi via multiple, gentle, washes with 1mL of phosphate buffered saline (PBS). Visual checks were performed to ensure A549s were intact, and planktonic NTHi were removed. Wells were then treated with 250µL of 0.25% Trypsin-EDTA to dislodge adherent bacteria (5min 37°C) before serial dilution and drop plating on columbia-blood agar (CBA) plates to enumerate bacterial loads. Results represent triplicate values of biological duplicates. The percentage adherence was calculated from the CFU in the inoculum.

Invasion assays were identical to the adherence assay with the following extra steps following removal of adhered bacteria: extracellular bacteria were killed via treatment with 100µg/mL gentamicin in RPMI containing 10% (v/v) FCS for 1h at 37°C. Effectiveness of gentamicin treatment was assessed by plating supernatant following treatment, with no bacterial growth evident. Wells were then treated with 250µL 0.2% (v/v) Saponin to lyse A549s (releasing intracellular bacteria). Visual checks were made to confirm cell lysis. Surviving intracellular NTHi were enumerated via serial dilution and drop plating as per adherence assays. Results represent triplicate values of biological duplicates. The percentage invasion was calculated from the CFU in the inoculum.

### Settling Assay

NTHi were grown in sBHI to an OD_600_ of 1.0. 3mL of OD_600_ 1.0 cells were resuspended in PBS, mixed thoroughly and then split into triplicate cuvettes per variant. Samples were monitored for 4 hours by measuring OD_600_. Values were expressed as % of initial reading.

### Adherence assays with ECM components

Flat bottom 96-well tissue culture treated plates (Falcon) were coated with vitronectin, laminin or fibronectin (all Sigma-Aldrich), according to manufacturer protocols. Briefly, working solutions of vitronectin (15 µg/ml) and laminin (6 µg/ml) were prepared in 1X DPBS, whereas fibronectin was reconstituted in water (15 µg/ml). From the working stock of vitronectin, 100 µl was added per well of 96 well plate and incubated at 37°C for 2 hours followed by overnight storage at 4°C. For laminin and fibronectin, 100 µl of working solutions was added to each well on the day of the assay, followed by immediate removal of the solution. Wells were air dried for 45 min and washed twice with 1X DPBS prior to the assay. Bacterial inoculum was prepared from log phase cultures of NTHi grown in sBHI and added at a density of 5×10^6^ CFU/well prepared in 1X DPBS to wells coated with individual ECM component. After incubation at 37°C and 5% CO_2_ for 1 hour, the supernatant was removed, and wells were washed 4 times with 1X DPBS to remove any non-adherent bacteria. Adherent bacteria were collected in 100 µl 1X DPBS with vigorous pipetting and scraping of the wells. Dilutions of the collected sample as well as the inoculum were plated on chocolate agar. The percentage adherence was calculated from the CFU in the inoculum.

### Biofilm imaging and analysis

Biofilms were formed by NTHi cultured within chambers of eight-well-chambered coverglass slides (Thermo Scientific, Waltham, MA) as described previously (33). Briefly, biofilms were formed by NTHi cultured within chambers of eight-well-chambered coverglass slides (Thermo Scientific, Waltham, MA) using mid-log-phase NTHi cultures. Bacteria were inoculated at 4 × 10^4^ CFU in 200-μl final volume per well and incubated at 37°C with 5% CO_2_ for 24 h, with the growth medium replaced after 16 h. To visualize, biofilms were stained with LIVE/DEAD BacLight stain (Life Technologies) and fixed overnight in fixative (1.6% paraformaldehyde, 2.5% glutaraldehyde, and 4% acetic acid in 0.1 M phosphate buffer, pH 7.4). Fixative was replaced with saline before imaging with a Zeiss 980 Meta-laser scanning confocal microscope. Images were rendered with Zeiss Zen software. Z-stack images were analyzed by COMSTAT2 (34) to determine biomass (μm3/μm2), average thickness (μm), and roughness (Ra).

### Phylogenetic tree

The 16s rRNA sequence of fully annotated *H. influenzae* genomes available in NCBI GenBank were aligned using CLUSTAL OMEGA (1.2.4). **Supplementary Table 1** contains full details of strains, genes and data used.

**Table 1.** Fragment Length Analysis of (A) Invasive and (B) Carriage Collections screened for *lav* SSR tract length using Lav_F and Lav_R. The results shown indicate whether the *lav* gene was ON (>70% ON; green), OFF (>70% OFF; red), mixed ON and OFF (orange), or if there was no gene (Grey) as we could not amplify a PCR product.

| **A) Invasive Collection** |  |  |  |  |
| --- | --- | --- | --- | --- |
| **OFF** | **ON** | **Mixed** | **No gene** | **Total** |
| 49 | 1 | 0 | 22 | **72** |
| 68.06 | 1.39 | 0.00 | 30.56 |  |
| Gene presence: 69.4% (50/72) |  |  |  |  |
| **B) Carriage Collection** |  |  |  |  |
| **OFF** | **ON** | **Mixed** | **No gene** | **Total** |
| 14 | 2 | 0 | 0 | **16** |
| 87.5% | 12.5% | 0% | 0% |  |
| Gene presence: 100% (16/16) |  |  |  |  |

### Statistical Analysis

Graphs and statistics were generated via GraphPad Prism 5.0 (GraphPad Software, La Jolla, California). Error bars represent standard deviation from mean values. A one-way ANOVA was used to compare samples: p values of <0.05 (considered significant) represented by *, p value of <0.001 indicated by **, p value of <0.001 indicated by ***. Groups were considered not significantly different if p > 0.05 (no *).

## Results

### Lav expression is phase-variable in NTHi due to changes in length an SSR tract in the open-reading frame

In order to study Lav function during colonisation and disease, we used prototype NTHi strain 86-028NP (35) that encoded *lav* (NTHI0585) and enriched populations of bacteria via single colony screening using fluorescent PCR (**Figure 1A**) for GCAA_(n)_ SSR tract lengths corresponding to all three possible reading frames. This resulted in three isogenic populations enriched for tracts containing 21 GCAA repeats (21r), 22 GCAA repeats (22r), and 23 GCAA repeats (23r) (**Figure 1B**). Analysis of *lav in silico* using the genome annotation from strain 86-028NP (Genbank accession number CP000057) determined that *lav* containing 21 GCAA_(n)_ repeats would be in-frame and ON (expressed), and those populations where the GCAA_(n)_ tract was 22 or 23 repeats would be out-of-frame and OFF (not expressed), due to premature transcriptional termination at stop codons in these two alternate reading frames. We also cloned and over-expressed the predicted passenger domain of Lav from 86-028NP (Lav-bind protein), based on previous analysis (22) to raise antisera. Western blots using this antisera and the three enriched populations confirmed our prediction that 21 repeats was ON (21r *lav* ON), and that 22 and 23 repeats were OFF (22r *lav* OFF and 23r *lav* OFF, respectively), as we could only detect the Lav protein in the 21r population (**Figure 1C**).

**Figure 1.**
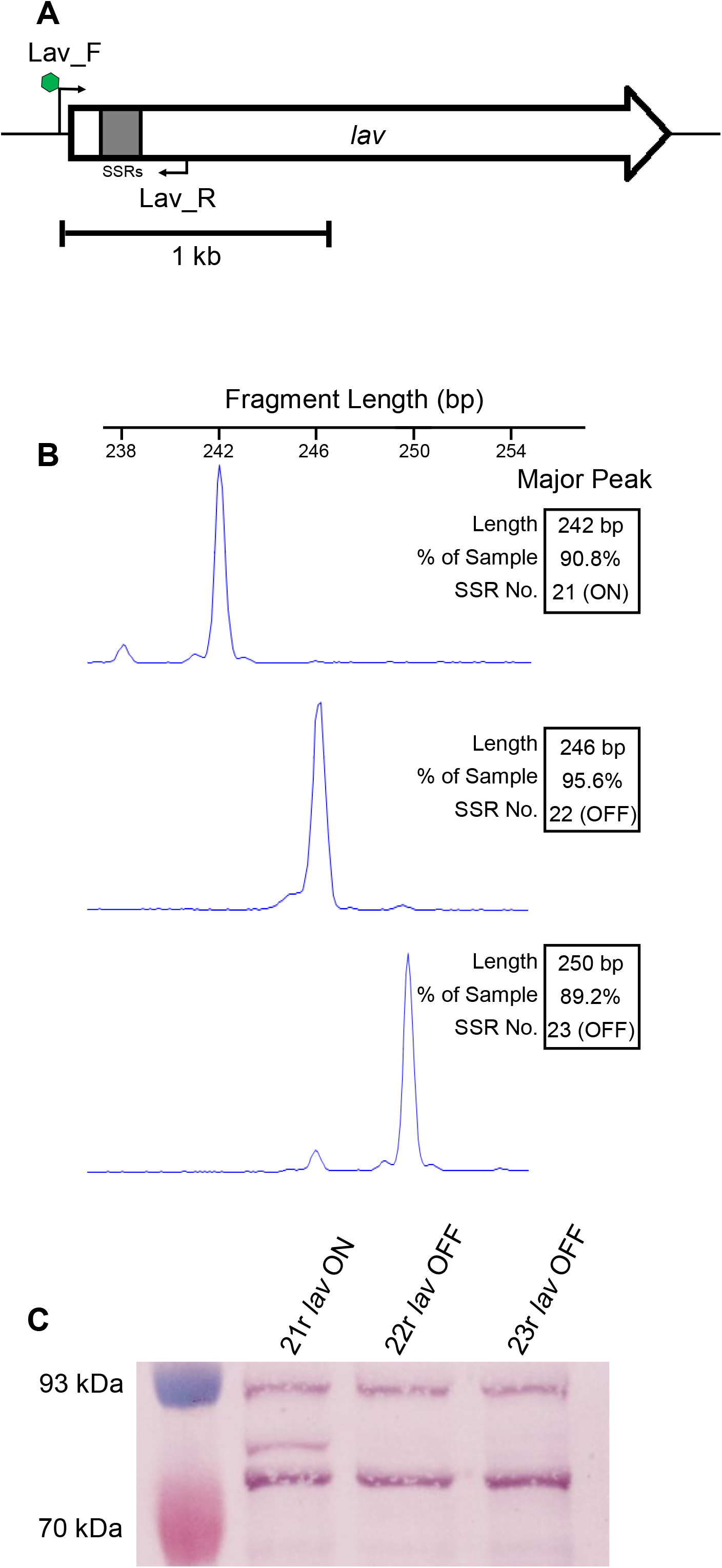
Phase variation of the *lav* gene. **(A)**. The 2.2kb *lav* gene, with 5’GCAA_(n)_ simple sequence repeat (SSR) tract in grey. A fluorescently (FAM) labelled forward primer (Lav_F; FAM indicated by green hexagon) binds upstream of the SSR tract. The reverse primer (Lav_R) binds downstream of the SSR tract. (**B)** fragment analysis traces of enriched variants for three consecutive GCAA_(n)_ repeat tract lengths (21r, 22r, 23r). (**C)**. Western Blot using whole-cell lysates of 86-028NP isogenic strains enriched in (B) with the *lav* gene containing an SSR number of 21, 22 or 23 repeats. An SSR tract number of 21 (21r) puts the gene in-frame, and ON, indicated by presence of the Lav protein detected using anti-Lav antisera. The 22r and 23r populations have the *lav* gene out-of-frame and OFF, with no Lav detected in cell lysates of these strains.

### The lav gene is switched OFF in NTHi isolates during both colonisation and invasive infection

We previously examined two collections of NTHi isolates for the expression of multiple lipooligosaccharide (LOS) biosynthetic enzymes (36), demonstrating that certain enzymes were selected for during invasive disease. We sought to further utilize these two collections to determine if phase variation of the *lav* gene occurred during colonisation and invasive disease. These two collections comprised carriage isolates, the ORChID collection (26, 27, 37), and a collection of invasive NTHi isolates (28). Fluorescently labelled PCR of the GCAA_(n)_ repeat tract of the *lav* gene (**Figure 1A**) was used to determine the ratio of each tract length present in the bacterial population, and calculate the percentage ON/OFF ratio of that population. Analysis of 16 isolates from our carriage collection showed that *lav* was present in all strains, and predominantly OFF in these isolates (14/16; 87.5%) (**Table 1**). Analysis of our invasive collection determined that the *lav* gene was present in ∼69% of the strains (**Table 1**), and where present, was also predominantly OFF (present in 50/72 isolates; of the 50 isolates encoding a *lav* gene, the gene is OFF in 49/50; 98%). This indicates that expression of *lav* may not be required during either colonisation or invasive infection, or there is a direct selection against expression of the Lav protein during both phenotypic states.

### Lav expression results in increased host cell adherence, but not invasion

In order to determine if Lav expression was required for an aspect of NTHi induced disease other than nasopharyngeal colonisation or invasive infection, we investigated the broad role of Lav during adherence to and invasion of the A549 human cell line, isolated from the lower human airway, using our ON/OFF enriched populations. These assays demonstrated that the Lav protein has a role in adherence to host cells, as 21r *lav* ON showed a significantly greater percentage of adherence than both 22r and 23r *lav* OFF variants (**Figure 2A**, p < 0.05). However, there was no significant difference in the ability of ON and OFF variants to invade these same cells (**Figure 2A**). The CFU/well and MOI values for both adherence and invasion assays are presented in **Supplementary Figure 2**. We also found that Lav expression is not required for inter-bacterial adherence, as there was no difference in the rate of settling as determined by using the optical density of a static culture of each of our enriched variants over 4 hours (**Figure 2B**).

**Figure 2.**
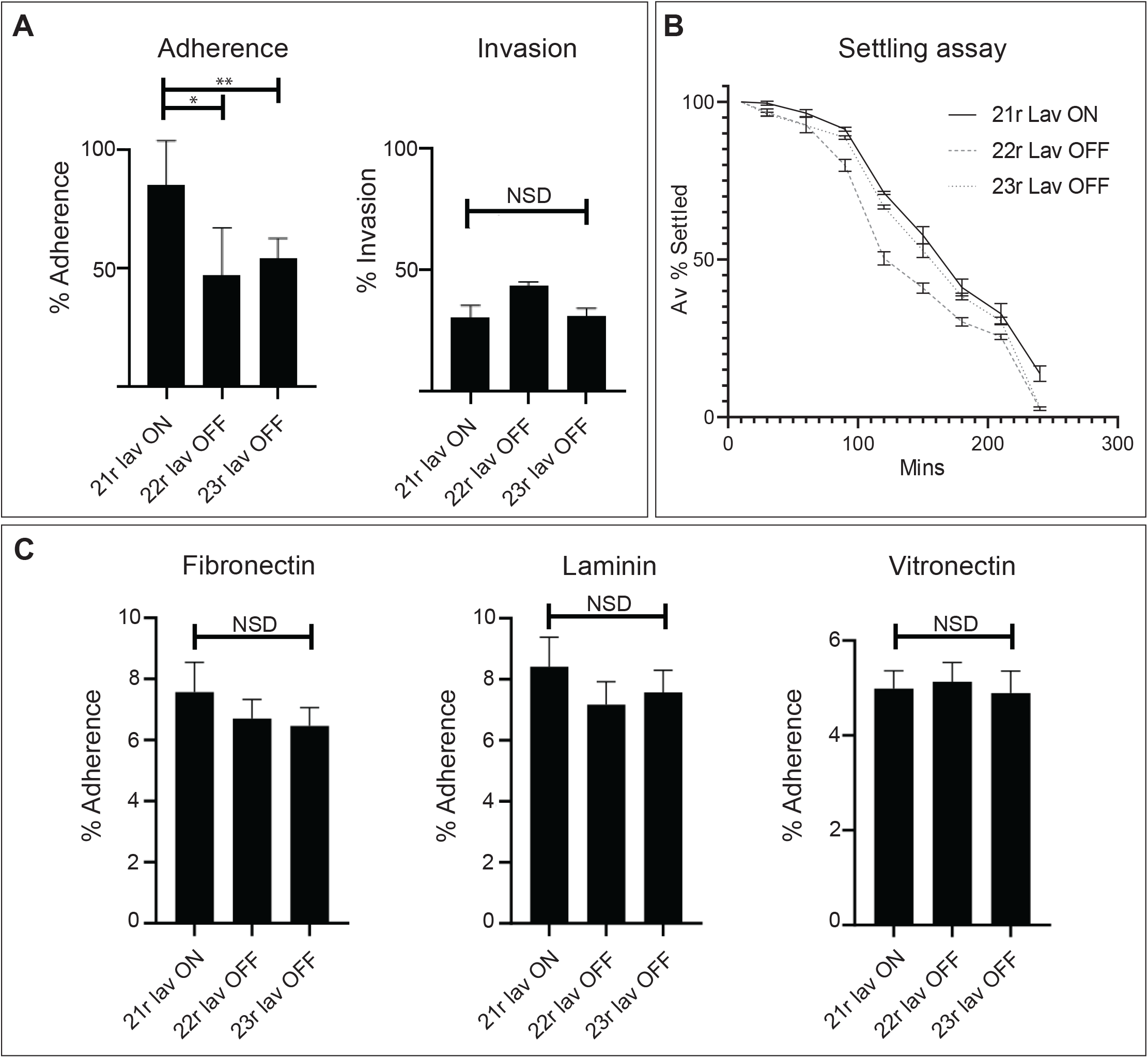
Impact of Lav expression on adherence and invasion. Impact of Lav expression on (**A**) adherence to and (**B**) invasion of the human A549 cell line was evaluated with Lav ON/OFF variants in NTHi strain 86-028NP, which expresses the Lav 1.2 variant. (**C**) Auto-aggregation was investigated by monitoring the OD_600_ of static cultures of our ON/OFF variants. **(D)** Adherence of NTHi *lav* variants to ECM proteins Fibronectin, Laminin, and Vitronectin. Statistical analysis was carried out by one way ANOVA. Error bars represent standard deviation from mean values; p value of <0.05 represented by *; p value of <0.001 indicated by ***; NSD = no significant difference between any of strains.

### Lav is not required for adherence to ECM components

Epithelial cells of the human respiratory tract produce multiple extracellular matrix (ECM) components (38-40). Since *lav* ON/OFF status affected the ability of NTHi to adhere to lung epithelial cells (**Figure 2A**), we tested adherence of the *lav* variants to the ECM components laminin, fibronectin and vitronectin for 1 hour. There was no significant difference observed between the percent adherence of the variants to laminin, fibronectin or vitronectin (**Figure 2C**), indicating that Lav is not involved in adherence of NTHi to these ECM components, but may instead be required to adhere specifically to receptor(s) only present on host cells.

### Lav expression results in biofilms with greater biomass and thickness

To determine if Lav phase variation resulted in differences in biofilm formation, a key feature of NTHi pathology, biofilms of our enriched 21r *lav* ON, 22r *lav* OFF, and 23r *lav* OFF variants were grown for 24 hours. Biofilms formed by the 21r *lav* ON variant exhibited significantly greater biomass and average thickness compared to variants that did not express Lav (22r and 23r *lav* OFF; **Figure 3A, 3B**). Biofilms formed by 22r *lav* OFF tended to have an architecture that was rougher in comparison to either of the other variants, likely due to the more dispersed nature of these biofilms, but the roughness of all three variants was statistically similar (**Figure 3C**). Based on gross biofilm abundance and microscopic analysis (**Figure 3D**), NTHi that expressed Lav (21r *lav* ON) formed significantly larger biofilms overall.

**Figure 3.**
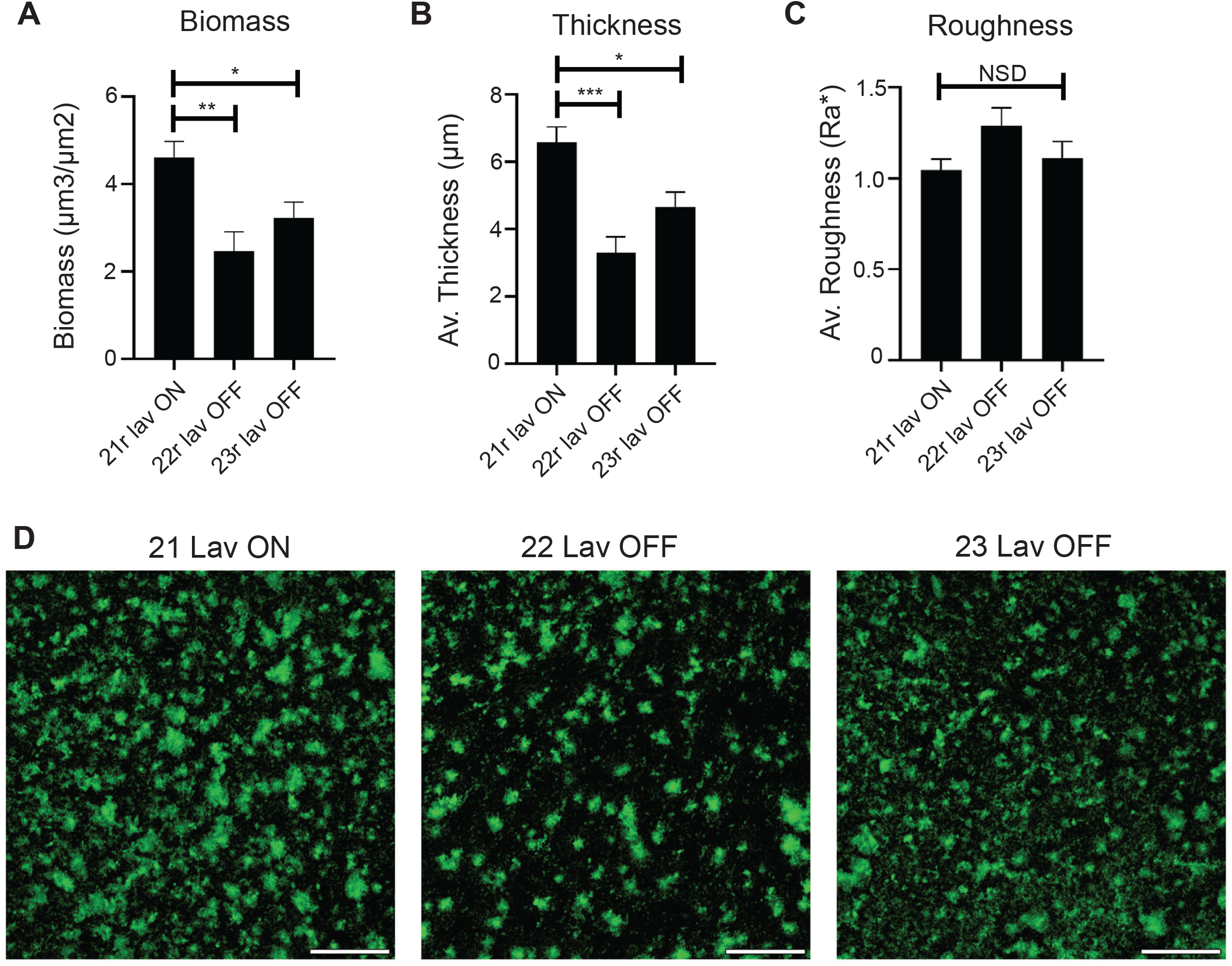
Impact of Lav expression on biofilm formation. **(A)** Biomass, **(B)** average thickness, and **(C)** roughness of biofilms grown for 24 h. Biofilms formed by the Lav expressing variant were of significantly greater biomass and thickness than those formed by the Lav non-expressing variants. Biofilms were analyzed by COMSTAT2, and values are shown as mean ± standard error of the mean. Statistical analysis was carried out by one way ANOVA. Error bars represent standard deviation from mean values; p value of <0.05 (considered significant) represented by *, p value of <0.01 indicated by **; p value of <0.001 indicated by ***; NSD = no significant difference between any of the strains. **(D)** Representative low-magnification images of biofilm density and distribution. Biofilms formed by 21r *lav* ON appeared denser with more and larger tower like structures compared to 22r *lav* OFF. 23r *lav* OFF formed biofilms with an intermediate distribution and smaller tower like structures. Bacteria are shown in green. Scale bar = 500 µm.

### *lav* distribution and conservation in *Haemophilus spp*

Previous studies demonstrated a broad distribution of *lav*, and multiple allelic variants of the Lav passenger domain in *Haemophilus* spp. (21, 22). However, there was no consistent naming of these variants in *Haemophilus* spp., nor a thorough analysis of the distribution or variability present. Therefore, we examined all fully annotated *Haemophilus* spp. genomes available in NCBI GenBank. There were 73 fully annotated *H. influenzae* genomes available at the time of this investigation. Of those 73, 47 were NTHi and the remainder were either typeable (serotypes a-f) or the serotype was undetermined. The *lav* gene was present in 29/47 NTHi genomes (∼62% gene presence), very similar to that observed in our invasive collection (69% presence). Interestingly, *lav* was absent in all strains annotated as serotype b-f, but was present in all strains (4) annotated as *H. influenzae* serotype a. A *lav* homologue, named *las*, was present in all 11 available genomes of *H. influenzae* biogroup *aegyptius* (7 fully annotated, plus 4 available genomes) (**Figure 4)**.

**Figure 4.**
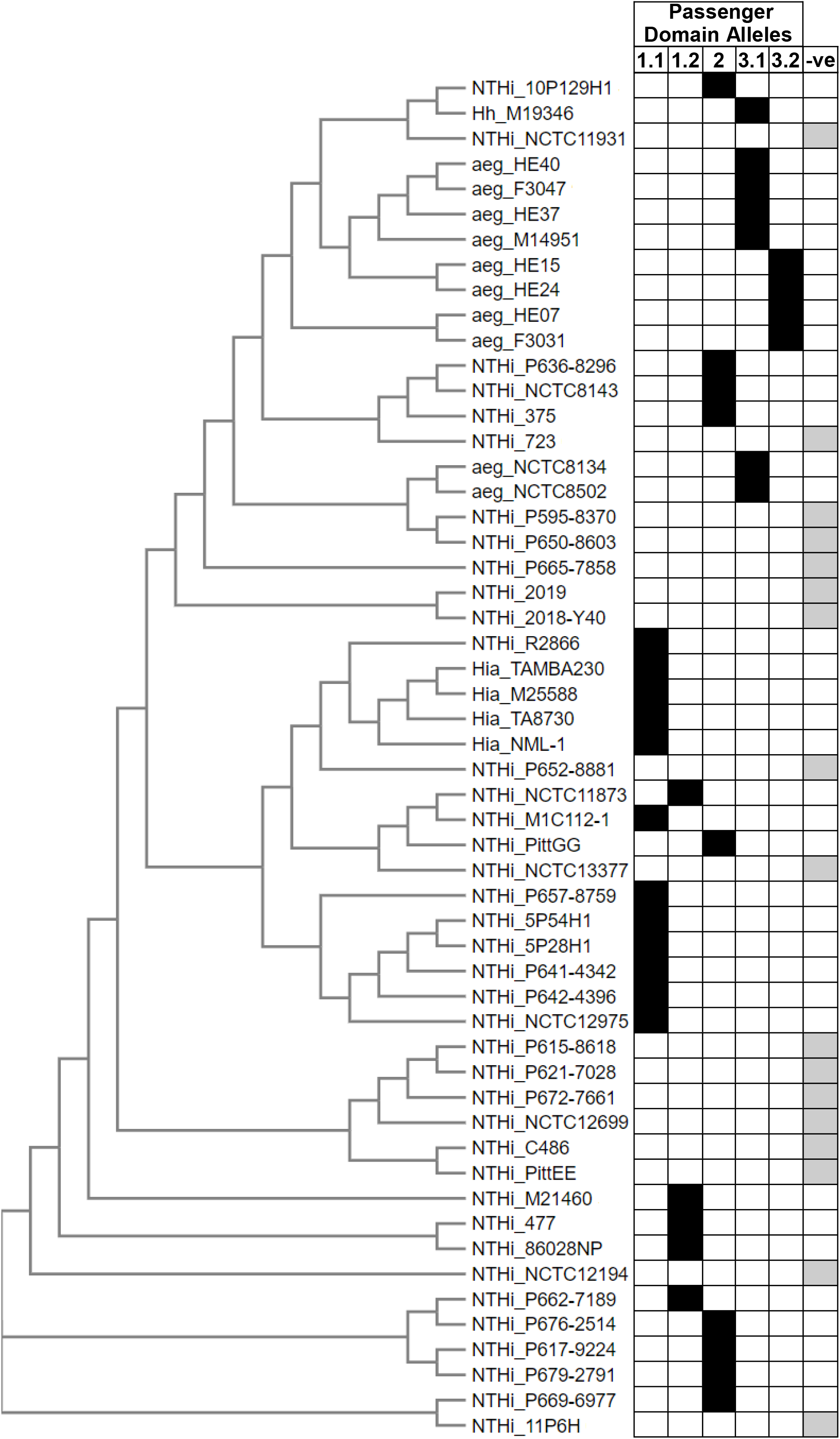
Distribution of *lav* gene in *Haemophilus spp*. Phylogenetic tree generated by CLUSTAL OMEGA (1.2.4) using 16s rRNA gene sequences from NTHi and *H. influenzae* serotype a GenBank entries. For sequences see **Supplementary Table 1**. Prefix describes species: NTHi – Non-Typeable *Haemophilus influenzae*; Hia – *Haemophilus influenzae serotype a*; aeg – *Haemophilus influenzae* biogroup *aegyptius*; Hh – *Haemophilus haemolyticus*. Suffix indicates strain name. i.e. “NTHi_86028NP” is Non-Typeable *Haemophilus influenzae* strain 86-028NP. Included in the figure is the Lav passenger domain allele form (**1.1, 1.2, 2, 3.1, 3.2**) and a negative column (**-ve**) to show genomes that did not contain the *lav* gene.

Further, we carried out detailed sequence analysis of all *lav* genes from these 73 *Haemophilus* spp. genomes, as previous work in *Neisseria* spp. had identified a number of allelic variants (21). Passenger domain variants 1 and 2 (previously named AutB1 and AutB2 (21)) were found exclusively in strains annotated as NTHi, with an approximate 60:40 split (59% and 41%, respectively). There appears to be a lineage distribution of these variants, with closely related strains containing the same passenger domain allele (**Figure 4**). Alignment of the sequences of the Lav passenger domain (**Figure 5**) showed that they were more diverse than previously described (21). Analysis of variant 1 showed two sub-variants present, with only 73.06% identity, and which we propose to name variants 1.1 and 1.2 (alignment in **Supplementary Figure 3**). With the exception of one *H. haemolyticus* strain, Lav passenger domain variant 3 (previously named AutB3 (21)) was found exclusively in *H. influenzae* biogroup *aegyptius* strains. Our sequence analysis of variant 3 also showed two distinct sub-variants, showing only 51.26% identity, and therefore we have also proposed delineating variant 3 into variants 3.1 and 3.2 (alignment in **Supplementary Figure 3**).

**Figure 5.**
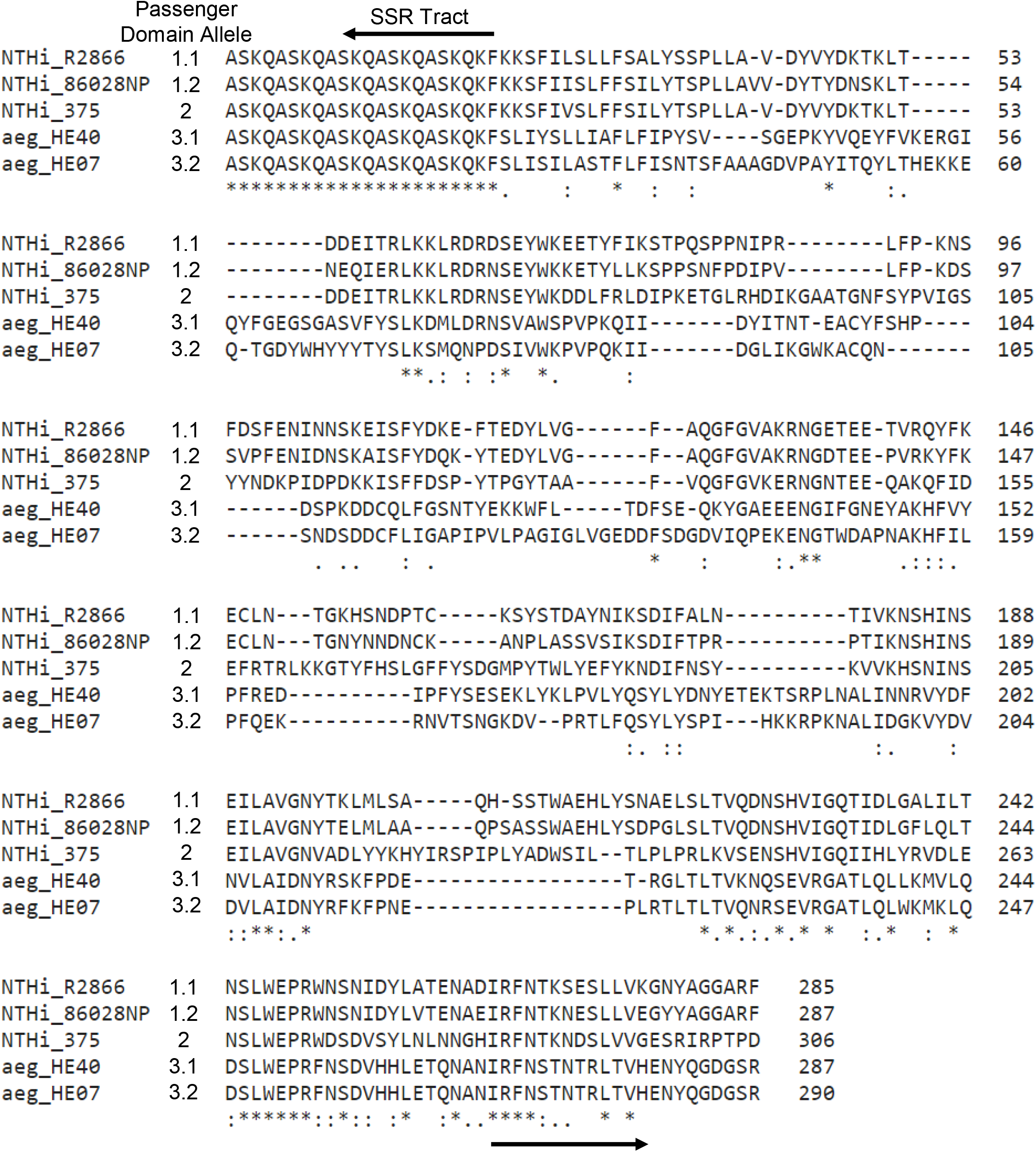
Alignment of the Lav Passenger Domain Alleles. CLUSTAL OMEGA (1.2.4) was used to align the distinct Lav passenger domain allele forms using representative amino acid sequences from five strains – R2866 (1.1), 86-028NP (1.2), 375 (2), HE40 (3.1) and HE07 (3.2).

## Discussion

Surface exposed NTHi phase-variable autotransporters are important virulence determinants (41). A Lav homologue, named AutB, was shown to be highly diverse, encoding multiple allelic variants of the functional passenger domain (21), with AutB important for biofilm formation in *N. meningitidis*. Previous work demonstrated that the *lav* gene in NTHi phase-varied OFF during repeated infection (22). Therefore, we aimed to determine the phenotypic role of Lav in NTHi, and to rationalise the prevalence and diversity of Lav in *Haemophilus* spp.

Analysis of our carriage and invasive NTHi collections (36) revealed *lav* to be present in ∼69% (50/72) of strains in our invasive collection (**Table 1A**), but in every strain in our carriage collection (**Table 1B**). Therefore, it appears that the invasive collection is representative of all NTHi strains, with *lav* found in ∼62% of fully annotated NTHi genomes (**Figure 4**). The high proportion of Lav observed in our carriage collection is likely an artefact of a small (16 isolates) sample size. Analysis of our invasive and carriage collections showed that *lav* is predominantly phase-varied OFF in NTHi colonising the nasopharynx (carriage collection; 87.5%) and during invasive infection (where present, *lav* is OFF in 49/50 isolates, equating to 98%), suggesting that Lav is either not required or directly selected against during both colonisation and invasive infection, or expressed in a distinct niche. Our invasive collection also represents just a ‘snap-shot’ of the exact phenotypic state at a particular time point during invasive infection, i.e., when treatment is required, which is likely to be during the later stages of disease. Previous work has shown that the *lav* gene switches OFF over subsequent episodes of infection (25), but can rapidly change expression over short periods (24). As we have found it to be OFF in the majority of both carriage and invasive isolates, it is possible that selection for the OFF state is due to negative selection from immune detection/pressure. Immune selection against multiple outer-membrane proteins has been reported in *N. meningitidis* (42), with gene expression phase-varying from ON to OFF during persistent carriage. Similar work with an additional phase-variable autotransporter in NTHi, Hia, showed Hia phase-varies OFF during opsonophagocytic killing (17). Our phenotypic findings regarding biofilm formation are also in agreement with previous work involving the Lav homologue AutB in *Neisseria meningitidis* (21). It therefore appears that Lav/AutB play a similar role in both species. The establishment of bacterial biofilms is critical during colonization and disease. Biofilms help bacteria adhere to the mucosal surfaces, and provide increased resistance to host defences and antimicrobials (43-45). Thus, expression of Lav might provide a selective advantage to NTHi during initial colonisation and establishment at the mucosal surface. Once established, factors such as immune pressures or microenvironmental conditions may select against the Lav expressing subpopulation as observed in the persistently colonized or invasive isolates assessed.

Our phenotypic analysis demonstrated that Lav has a role in adherence to, but not for invasion of, human A549 lung cells (**Figure 2A**), and that Lav expression does not play a role in adherence to ECM proteins (**Figure 2C)**. This indicates that there is a state in which Lav expression is beneficial, perhaps during the early stages of colonisation or during progression from the upper to the lower respiratory tract, or whilst establishing invasive/systemic infection. Another possibility is that Lav interacts with a specific receptor present on human cells, and does not directly bind to the ECM proteins fibronectin, laminin, and vitronectin. The NTHi autotransporter Hia has been shown to bind to human specific glycans rather than proteins as high affinity receptors (46).

Understanding the prevalence and conservation of Lav is key for determining its suitability for use in a rationally designed subunit vaccine against NTHi. Our analysis determined that the *lav* gene is found in ∼62% of *Haemophilus* spp. Intriguingly, there was no *lav* gene, or close homologue, in any strain annotated as *H. influenzae* serotype b-f. The presence of Lav in *H. influenzae* serotype a only may suggest that this protein is an important virulence factor in these strains, although the small number of sequences publicly available for analysis means this hypothesis will require further investigation.

Our investigation of the diversity of the Lav passenger domains present in *Haemophilus* spp. showed that there are five different allelic variants of the Lav passenger domain (the functional extracellular region) present in *Haemophilus* spp. Our detailed sequence analysis showed that both variants 1 and 3 can be further divided into two separate allelic variants – 1.1 and 1.2, and 3.1 and 3.2 (**Figure 5**). Variants 3.1 and 3.2, previously annotated as Las (22, 23), are found exclusively in *H. influenzae* biogroup *aegyptius* isolates.

*H. influenzae* biogroup *aegyptius* strains cause the invasive disease Brazilian purpuric fever (BPF), a meningitis-like disease with high fatality (47). The ubiquitous presence of Lav variants 3.1 and 3.2 in *H. influenzae* biogroup *aegyptius* isolates supports the idea that these particular variants contribute to the development of BPF, although it has previously been reported that no single factor is required for BPF (47). It has also previously been reported that *las* expression is highly variable during an animal model of BPF (24), with expression of the gene shown to decrease (switch OFF) after 24 hours, then increase (switch ON) at the 48 hour time point post infection, demonstrating complex regulation of Lav/Las occurs during disease. In summary, our analysis has shown that there are five unique variants of the Lav passenger domain encoded by *Haemophilus* spp., and there is a distinct distribution between serotype/species. Future investigation into the functional differences between passenger domain variants is needed to determine if these variants have different functions.

It is important to understand the role of bacterial surface factors like Lav in order to understand NTHi-mediated diseases, and to develop effective vaccines and treatments. Our work has determined that expression of a particular Lav variant (1.2 in strain 86-028NP) results in greater host cell adherence and biofilm formation, and demonstrated that the *lav* gene is present as five different allelic variants in *Haemophilus* spp. As Lav is present in ∼60% of NTHi strains, understanding the role of all variants is key to understanding NTHi disease, and further work is required to assess if Lav can form part of a multi-subunit, rationally designed vaccine against NTHi.

## Acknowledgements

We thank the Department of Cell Biology, Neurobiology and Anatomy at MCW for the use of their Zeiss LSM980 confocal microscope. This work was supported by an Australian Research Council (ARC) Discovery Project grant (DP180100976) to J.M.A. an Australian National Health and Medical Research Council (NHMRC) Principal Research Fellowship (1138466) to M.P.J, and a National Institutes of Health (NIH) grant (R21-DC016709) to K.L.B. The ORChID study was supported by a NHMRC project grant (GNT615700) and a program grant from the Children’s Health Foundation Queensland (5006). Publication costs of this work were supported by a generous donation from the Bourne Foundation, Melbourne, Australia.

## References

1. Murphy TF, Faden H, Bakaletz LO, Kyd JM, Forsgren A, Campos J, Virji M, Pelton SI. 2009. Nontypeable Haemophilus influenzae as a pathogen in children. Pediatr Infect Dis J 28:43–8.

2. Sethi S, Murphy TF. 2008. Infection in the pathogenesis and course of chronic obstructive pulmonary disease. N Engl J Med 359:2355–65.

3. Van Eldere J, Slack MP, Ladhani S, Cripps AW. 2014. Non-typeable Haemophilus influenzae, an under-recognised pathogen. Lancet Infect Dis 14:1281–92.

4. Johnson RH. 1988. Community-acquired pneumonia: etiology, diagnosis, and treatment. Clin Ther 10:568–73.

5. Ladhani S, Slack MP, Heath PT, von Gottberg A, Chandra M, Ramsay ME. 2010. Invasive Haemophilus influenzae Disease, Europe, 1996-2006. Emerg Infect Dis 16:455–63.

6. Collins S, Vickers A, Ladhani SN, Flynn S, Platt S, Ramsay ME, Litt DJ, Slack MP. 2016. Clinical and Molecular Epidemiology of Childhood Invasive Nontypeable Haemophilus influenzae Disease in England and Wales. Pediatr Infect Dis J 35:e76–84.

7. Slack MPE, Cripps AW, Grimwood K, Mackenzie GA, Ulanova M. 2021. Invasive Haemophilus influenzae Infections after 3 Decades of Hib Protein Conjugate Vaccine Use. Clin Microbiol Rev 34:e0002821.

8. Atkinson CT, Kunde DA, Tristram SG. 2017. Expression of acquired macrolide resistance genes in Haemophilus influenzae. J Antimicrob Chemother 72:3298–3301.

9. Phillips ZN, Tram G, Seib KL, Atack JM. 2019. Phase-variable bacterial loci: how bacteria gamble to maximise fitness in changing environments. Biochem Soc Trans 47:1131–1141.

10. Phillips ZN, Husna AU, Jennings MP, Seib KL, Atack JM. 2019. Phasevarions of bacterial pathogens - phase-variable epigenetic regulators evolving from restriction-modification systems. Microbiology doi:10.1099/mic.0.000805.

11. Moxon R, Bayliss C, Hood D. 2006. Bacterial contingency loci: the role of simple sequence DNA repeats in bacterial adaptation. Annu Rev Genet 40:307–33.

12. Jeffrey N. Weuser DJM, Peter D. Butler, Alf A. Lindberg, E. Richard Moxon. 1990. Characterization of repetitive sequences controlling phase variation of Haemophilus influenzae Lipopolysaccharide. Journal of Bacteriology:3304 –3309.

13. Elango D, Schulz BL. 2020. Phase-Variable Glycosylation in Nontypeable Haemophilus influenzae. J Proteome Res 19:464–476.

14. Meuskens I, Saragliadis A, Leo JC, Linke D. 2019. Type V Secretion Systems: An Overview of Passenger Domain Functions. Frontiers in Microbiology 10.

15. Battaglioli EJ, Goh KGK, Atruktsang TS, Schwartz K, Schembri MA, Welch RA. 2018. Identification and Characterization of a Phase-Variable Element That Regulates the Autotransporter UpaE in Uropathogenic Escherichia coli. mBio 9.

16. Spahich NA, Hood DW, Moxon ER, St Geme JW, 3rd. 2012. Inactivation of Haemophilus influenzae lipopolysaccharide biosynthesis genes interferes with outer membrane localization of the hap autotransporter. J Bacteriol 194:1815–22.

17. Atack JM, Winter LE, Jurcisek JA, Bakaletz LO, Barenkamp SJ, Jennings MP. 2015. Selection and counter-selection of Hia expression reveals a key role for phase-variable expression of this adhesin in infection caused by non-typeable Haemophilus influenzae. J Infect Dis 212:645–53.

18. Serruto D, Spadafina T, Ciucchi L, Lewis LA, Ram S, Tontini M, Santini L, Biolchi A, Seib KL, Giuliani MM, Donnelly JJ, Berti F, Savino S, Scarselli M, Costantino P, Kroll JS, O’Dwyer C, Qiu J, Plaut AG, Moxon R, Rappuoli R, Pizza M, Aricò B. 2010. Neisseria meningitidis GNA2132, a heparin-binding protein that induces protective immunity in humans. Proceedings of the National Academy of Sciences of the United States of America 107:3770–3775.

19. van Ulsen P, van Alphen L, ten Hove J, Fransen F, van der Ley P, Tommassen J. 2003. A Neisserial autotransporter NalP modulating the processing of other autotransporters. Mol Microbiol 50:1017–30.

20. Arenas J, Cano S, Nijland R, van Dongen V, Rutten L, van der Ende A, Tommassen J. 2015. The meningococcal autotransporter AutA is implicated in autoaggregation and biofilm formation. Environ Microbiol 17:1321–37.

21. Arenas J, Paganelli FL, Rodríguez-Castaño P, Cano-Crespo S, van der Ende A, van Putten JPM, Tommassen J. 2016. Expression of the Gene for Autotransporter AutB of Neisseria meningitidis Affects Biofilm Formation and Epithelial Transmigration. Frontiers in Cellular and Infection Microbiology 6.

22. Davis J SA, Hughes WR, Golomb M. 2001. Evolution of an autotransporter: domain shuffling and lateral transfer from pathogenic Haemophilus to Neisseria. J Bacteriol 2001;183(15):4626–4635..

23. Cury GCG, Pereira RFC, de Hollanda LM, Lancellotti M. 2015. Inflammatory response of Haemophilus influenzae biotype aegyptius causing Brazilian Purpuric Fever. Brazilian journal of microbiology : [publication of the Brazilian Society for Microbiology] 45:1449–1454.

24. Pereira RFC, Theizen TH, Machado D, Guarnieri JPO, Gomide GP, Hollanda LM, Lancellotti M. 2020. Analysis of potential virulence genes and competence to transformation in Haemophilus influenzae biotype aegyptius associated with Brazilian Purpuric Fever. Genet Mol Biol 44:e20200029.

25. Harrison A, Hardison RL, Fullen AR, Wallace RM, Gordon DM, White P, Jennings RN, Justice SS, Mason KM. 2020. Continuous Microevolution Accelerates Disease Progression during Sequential Episodes of Infection. Cell Rep 30:2978-2988.e3.

26. Lambert SB, Ware RS, Cook AL, Maguire FA, Whiley DM, Bialasiewicz S, Mackay IM, Wang D, Sloots TP, Nissen MD, Grimwood K. 2012. Observational Research in Childhood Infectious Diseases (ORChID): a dynamic birth cohort study. BMJ Open 2.

27. Sarna M, Lambert SB, Sloots TP, Whiley DM, Alsaleh A, Mhango L, Bialasiewicz S, Wang D, Nissen MD, Grimwood K, Ware RS. 2018. Viruses causing lower respiratory symptoms in young children: findings from the ORChID birth cohort. Thorax 73:969–979.

28. Staples M, Graham RMA, Jennison AV. 2017. Characterisation of invasive clinical Haemophilus influenzae isolates in Queensland, Australia using whole-genome sequencing. Epidemiol Infect 145:1727–1736.

29. Phillips ZN, Brizuela C, Jennison AV, Staples M, Grimwood K, Seib KL, Jennings MP, Atack JM. 2019. Analysis of invasive non-typeable Haemophilus influenzae isolates reveals a selection for the expression state of particular phase-variable lipooligosaccharide biosynthetic genes. Infect Immun 87:10.1128/iai.00093-19.

30. Tram G, Jen FE, Phillips ZN, Timms J, Husna AU, Jennings MP, Blackall PJ, Atack JM. 2021. Streptococcus suis Encodes Multiple Allelic Variants of a Phase-Variable Type III DNA Methyltransferase, ModS, That Control Distinct Phasevarions. mSphere 6:e00069–21.

31. Goyal M, Singh M, Ray P, Srinivasan R, Chakraborti A. 2015. Cellular interaction of nontypeable Haemophilus influenzae triggers cytotoxicity of infected type II alveolar cells via apoptosis. Pathogens and Disease 73:1–12.

32. Geme JWS, Falkow S. 1990. Haemophilus influenzae adheres to and enters cultured human epithelial cells. Infection and Immunity 58:4036–4044.

33. Jurcisek JA, Dickson AC, Bruggeman ME, Bakaletz LO. 2011. In vitro biofilm formation in an 8-well chamber slide. Journal of Visualized Experiments doi:10.3791/2481:pii: 2481. doi: 10.3791/2481.

34. Heydorn A, Nielsen AT, Hentzer M, Sternberg C, Givskov M, Ersbøll BK, Molin S. 2000. Quantification of biofilm structures by the novel computer program COMSTAT. Microbiology (Reading) 146 (Pt 10):2395–2407.

35. Harrison A, Dyer DW, Gillaspy A, Ray WC, Mungur R, Carson MB, Zhong H, Gipson J, Gipson M, Johnson LS, Lewis L, Bakaletz LO, Munson RS, Jr. 2005. Genomic sequence of an otitis media isolate of nontypeable Haemophilus influenzae: comparative study with H. influenzae serotype d, strain KW20. J Bacteriol 187:4627–36.

36. Phillips ZN, Brizuela C, Jennison AV, Staples M, Grimwood K, Seib KL, Jennings MP, Atack JM. 2019. Analysis of Invasive Nontypeable Haemophilus influenzae Isolates Reveals Selection for the Expression State of Particular Phase-Variable Lipooligosaccharide Biosynthetic Genes. Infect Immun 87.

37. Palmu AA, Ware RS, Lambert SB, Sarna M, Bialasiewicz S, Seib KL, Atack JM, Nissen MD, Grimwood K. 2019. Nasal swab bacteriology by PCR during the first 24-months of life: A prospective birth cohort study. Pediatr Pulmonol 54:289–296.

38. Coraux C, Roux J, Jolly T, Birembaut P. 2008. Epithelial cell-extracellular matrix interactions and stem cells in airway epithelial regeneration. Proc Am Thorac Soc 5:689–94.

39. Peters-Hall JR, Brown KJ, Pillai DK, Tomney A, Garvin LM, Wu X, Rose MC. 2015. Quantitative proteomics reveals an altered cystic fibrosis in vitro bronchial epithelial secretome. Am J Respir Cell Mol Biol 53:22–32.

40. Salazar-Peláez LM, Abraham T, Herrera AM, Correa MA, Ortega JE, Paré PD, Seow CY. 2015. Vitronectin expression in the airways of subjects with asthma and chronic obstructive pulmonary disease. PLoS One 10:e0119717.

41. Spahich NA, St Geme JW, 3rd. 2011. Structure and function of the Haemophilus influenzae autotransporters. Frontiers in cellular and infection microbiology 1:5–5.

42. Alamro M, Bidmos FA, Chan H, Oldfield NJ, Newton E, Bai X, Aidley J, Care R, Mattick C, Turner DPJ, Neal KR, Ala’Aldeen DAA, Feavers I, Borrow R, Bayliss CD. 2014. Phase Variation Mediates Reductions in Expression of Surface Proteins during Persistent Meningococcal Carriage. Infection and Immunity 82:2472–2484.

43. Yan J, Bassler BL. 2019. Surviving as a Community: Antibiotic Tolerance and Persistence in Bacterial Biofilms. Cell Host Microbe 26:15–21.

44. Murphy TF, Bakaletz LO, Smeesters PR. 2009. Microbial interactions in the respiratory tract. Pediatr Infect Dis J 28:S121–6.

45. Novotny LA, Brockman KL, Mokrzan EM, Jurcisek JA, Bakaletz LO. 2019. Biofilm biology and vaccine strategies for otitis media due to nontypeable Haemophilus influenzae. J Pediatr Infect Dis 14:69–77.

46. Atack JM, Day CJ, Poole J, Brockman KL, Timms JRL, Winter LE, Haselhorst T, Bakaletz LO, Barenkamp SJ, Jennings MP. 2020. The Nontypeable Haemophilus influenzae Major Adhesin Hia Is a Dual-Function Lectin That Binds to Human-Specific Respiratory Tract Sialic Acid Glycan Receptors. mBio 11.

47. Harrison LH, Simonsen V, Waldman EA. 2008. Emergence and disappearance of a virulent clone of Haemophilus influenzae biogroup aegyptius, cause of Brazilian purpuric fever. Clin Microbiol Rev 21:594–605.

